# Somite morphogenesis is required for axial blood vessel formation

**DOI:** 10.1101/2021.04.07.438831

**Authors:** Eric Paulissen, Nicholas J. Palmisano, Joshua Waxman, Benjamin L. Martin

## Abstract

Angioblasts that form the major axial blood vessels of the dorsal aorta and cardinal vein migrate towards the embryonic midline from distant lateral positions. Little is known about what controls the precise timing of angioblast migration and their final destination at the midline. Using zebrafish, we found that midline angioblast migration requires neighboring tissue rearrangements generated by somite morphogenesis. The somitic shape changes cause the adjacent notochord to separate from the underlying endoderm, creating a ventral midline cavity that provides a physical space for the angioblasts to migrate into. The anterior to posterior progression of midline angioblast migration is facilitated by retinoic acid induced anterior to posterior somite maturation and the subsequent progressive opening of the ventral midline cavity. Our work demonstrates a critical role for somite morphogenesis in organizing surrounding tissues to facilitate notochord positioning and angioblast migration, which is ultimately responsible for creating a functional cardiovascular system.

**Summary statement:** Retinoic acid induced somite morphogenesis generates a midline cavity that accommodates migrating angioblasts, which form the axial blood vessels.

## INTRODUCTION

Early organismal development relies on a variety of tissues that collectively organize into a functional body plan. In the trunk of the vertebrate embryo, many tissues are differentiating at once, and often interact with one another to influence their definitive structure (McMillen and Holley, 2015). The axial (notochord), paraxial (somites), and lateral (angioblasts and other tissues) mesoderm begins to be specified during gastrulation, and new cells are added to these tissues from posteriorly localized progenitor cells as they expand after gastrulation (Kimelman, 2016; Martin, 2016; Martin and Kimelman, 2012; Row et al., 2016). By the end of embryogenesis, the notochord is a prominent midline tissue of the trunk, and provides structural support to the body and signals that pattern adjacent tissues (Balmer et al., 2016; Glickman et al., 2003; Kimmel et al., 1995). Pairs of somites flank either side of the notochord and give rise to tissues that include the skeletal muscle, tendons, and bone (Tani et al., 2020). Ventral to the notochord and between the somites, the axial vasculature structures of the dorsal aorta and posterior cardinal vein distribute blood throughout the embryo (Hogan and Bautch, 2004; Isogai et al., 2001).

During somitogenesis, newly formed somites undergo a maturation event in which they change their morphology from a cuboidal shape to a chevron shape, and extend in the dorsal-ventral axis (Kimmel et al., 1995; Tlili et al., 2019). Many cellular rearrangements and mechanical stresses contribute to making the final definitive somite shape (Hollway et al., 2007; Leal et al., 2014; Tlili et al., 2019; Yin and Solnica-Krezel, 2007; Yin et al., 2018; Youn and Malacinski, 1981). At the same time that somites are changing shape, endothelial progenitors called angioblasts arise in the lateral plate mesoderm and migrate to the midline of the embryo to form the axial vasculature (Jin et al., 2005).

Interestingly, angioblast migration and new somite formation occur in an anterior to posterior progression, with a wave of angioblast migration happening slightly after the wave of somitogenesis (Jin et al., 2005; Kohli et al., 2013; Yabe and Takada, 2016). Despite these two events occurring in close temporal and physical proximity to one another, it is not clear how they influence one another during this developmental stage. While some evidence has shown somite-related defects on blood vessel patterning, much of this research focused on angiogenic sprouting or defects in arterial-venous specification, well after angioblast migration and somitogenesis is completed (Lawson et al., 2002; Shaw et al., 2006; Therapontos and Vargesson, 2010; Torres-Vázquez et al., 2004).

Loss of function mutation of the notochord specifying gene noto gene results in non-autonomous migration defects of angioblasts as they move to the midline. (Fouquet et al., 1997; Helker et al., 2015). In the absence of noto, the notochord adopts a somitic fate (Fouquet et al., 1997; Talbot et al., 1995). The requirement of notochord specification for angioblast migration suggested that the notochord may act to attract angioblasts to the midline. This model was further explored when a notochord-derived secreted factor named apela (also known as toddler/elabela) was discovered (Chng et al., 2013; Freyer et al., 2017; Helker et al., 2015; Pauli et al., 2014). This small peptide is expressed in the notochord and when mutated caused angioblast migration defects.

However, apela loss of function also causes defects in mesoderm and endoderm formation prior to notochord and angioblast specification, via gastrulation defects (Freyer et al., 2017; Norris et al., 2017; Pauli et al., 2014). Similarly, noto loss of function causes morphological changes that could interfere with angioblast migration, including a broad expansion of somite tissue near the developing blood vessels (Halpern et al., 1995). Thus, although it appears that mutations that affect notochord development can disrupt angioblast migration, the exact mechanism of this effect is not clear.

As new somites form during body axis extension, they secrete all-trans retinoic acid (RA). RA is a metabolic derivative of vitamin A, and a series of alcohol and aldehyde dehydrogenases convert Vitamin A into RA in specific locations of the embryo including newly formed somites (Duester, 2008). RA acts as a morphogen with broad roles during vertebrate embryogenesis. Some of these include the development of the limb, skeleton, hindbrain, and heart (Emoto et al., 2005; Heine et al., 1986; Lohnes et al., 1994; Mendelsohn et al., 1994; Niederreither et al., 2001; Sandell et al., 2007). Although involved in many processes, RA has a particular influence over the maturation of mesoderm and somitic tissue (Hamade et al., 2006; Janesick et al., 2018, 2014; Li et al., 2015). While RA is notably involved in ensuring the bilateral symmetry of the somites, it is ultimately dispensable for axis elongation in the zebrafish (Berenguer et al., 2018; Bernheim and Meilhac, 2020; Hamade et al., 2006; Kumar and Duester, 2014). Based on the timing of somite derived RA signaling activity relative to when angioblasts migrate to the midline, we speculated it may be involved in midline angioblast migration.

Here we show that RA mediated somite maturation is required for the proper formation of the axial vasculature through a non-autonomous role in inducing morphological changes in surrounding tissues. A process we call notochord-endoderm separation (NES) occurs prior to angioblast migration, wherein the dorsal translocation of the notochord during development leads to its separation from the endoderm along the dorsal-ventral axis to generate a transient cavity we refer to as the ventral midline cavity (VMC). The angioblasts migrate toward and eventually into the VMC after NES. The induction of NES and the VMC is somite maturation dependent, and a delay or failure in NES can cause systemic angioblast migration defects. This evidence places somite- maturation as a critical event that is required for NES and the development of the axial vasculature.

## RESULTS

### Retinoic acid promotes convergence of angioblasts to the midline and the formation of the axial vasculature

To determine if RA signaling impacts angioblast migration, we made time-lapse movies of tg(kdrl:GFP) embryos, which express GFP in angioblasts (Jin et al., 2005). Embryos were mounted at the 10 somite-stage and their migration was observed over a 3-hour time period. Wild-type angioblasts migrated to the midline in a manner consistent with previously described research (Figure 1A, Supplemental Movie 1) (Helker et al., 2015; Jin et al., 2005; Kohli et al., 2013). To determine what effect the loss of RA would have on the migrating angioblasts, we generated aldh1a2 mutants with the tg(kdrl:GFP) angioblast reporter in the background. Time-lapse imaging of these embryos shows defects in angioblast migration to the midline (Figure 1B, Supplemental Movie 2), which was also confirmed by in-situ hybridization (Figure S1). Three other methods of disrupting RA signaling caused the same phenotype. Loss of RA by treatment with N,N-diethylaminobenzaldehyde (DEAB) (an inhibitor of the Aldh family, including Aldh1a2 (Morgan et al., 2015)), the small molecule retinoic acid receptor (RAR) alpha and gamma inhibitor (BMS453), or expression of a dominant negative RARa using the Tg(hsp70l:EGFP-dnHsa.RARA) transgenic line (hereafter referred to as HS:dnRAR) all caused midline migration defects (Skvarca et al., 2019) (Figure S1). Angioblasts in these embryos show delayed migration with disorganized anterior to posterior processivity. To determine if addition of RA accelerated angioblast migration as well as anterior to posterior processivity, we compared the angioblast migration patterns of 10 somite stage tg(kdrl:GFP) embryos treated with either DMSO (vehicle) or 0.1 μM RA at the tailbud stage (Figure 1C,1D, respectively). Angioblasts in the DMSO treated embryos migrate normally, but the angioblasts treated with RA had prematurely initiated migration and arrived earlier to the midline during development (Figure 1C and Figure 1D, respectively, Supplemental Movie 3). To determine the rate at which angioblasts reach the midline, we quantified the percentage of GFP fluorescence at the midline in relation to the total amount of fluorescence for the both wild-type and aldh1a2 -/-. We defined angioblast fluorescence as reaching the midline if they are settled beneath the notochord. The aldh1a2 -/- embryos show very low percentage fluorescence at the midline, relative to total fluorescence, compared to wild-type embryos 4 hours after observation, when angioblast migration is largely complete (Figure 1E). On the other hand, fluorescence was higher at the midline in embryos treated with RA than in DMSO treated control embryos (Figure 1F). This shows that RA is necessary and sufficient for promoting angioblast migration to the midline. To determine at which stage RA is needed for angioblasts to migrate to the midline, we administered DEAB at different stages of development. Inhibition of RA at tailbud stage caused the angioblast migration defect, but treatment at the 5-somite stage resulted in normal migration (Figure 1H and 1I, respectively), thus establishing the critical developmental window for RA signaling regulation of midline angioblast migration.

**Figure 1.**
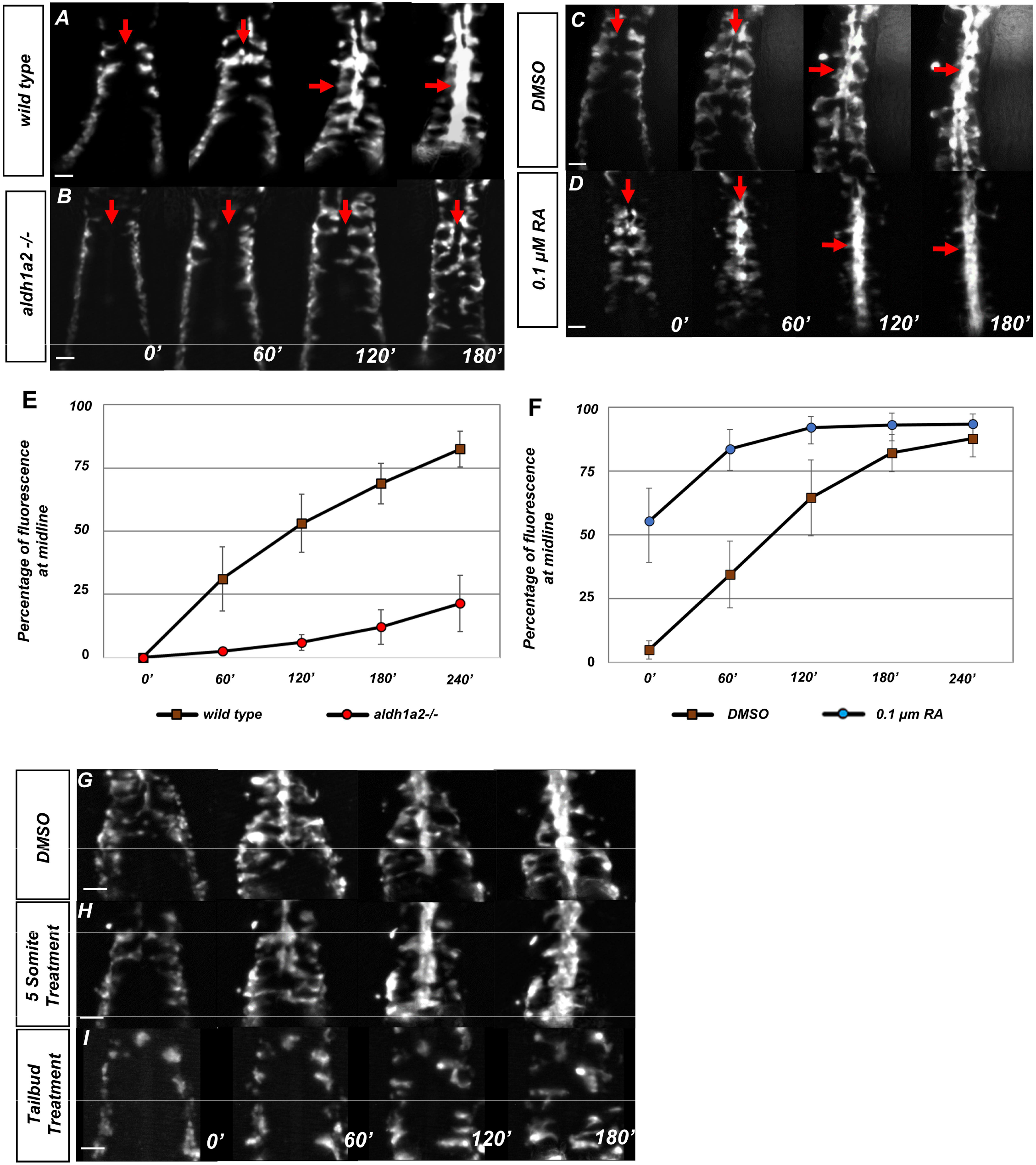
Retinoic acid is required prior to segmentation for angioblast migration. (A-D) Dorsal view of tg(kdrl:eGFP) embryos with representative images taken every 60 minutes over the course of 180 minutes, in (A) untreated (N=12, see Supplemental Movie 1), (B) aldh1a2 -/- mutants (N=12, see Supplemental Movie 2), (C) DMSO treated (N=10), or 0.1 μM RA treated embryos (N=8, see Supplemental Movie 3). The anterior of the embryo is at the top and red arrows indicate the midline. (E, F) Graph of fluorescence intensity as angioblasts arrive at the midline for (E) wild-type and aldh1a2 -/- and for (F) DMSO and 0.1 μM RA. Y-axis indicates percentage of fluorescent intensity at midline relative to the fluorescence intensity of whole embryo. Time is measured on the X-axis. N=8 for all conditions, error bars indicate standard deviation. (G-I) time-lapse images of tg(kdrl:eGFP) embryos treated with either (G) DMSO vehicle (n=10) or (H) 20 μM DEAB at the 5 somite stage (n=6), and (I) 20 μM DEAB at tailbud stage (n=14). Scale bars, 50 μm.

### Retinoic acid is required cell non-autonomously for angioblast migration

To determine if RA functions cell autonomously to induce angioblast migration, we utilized the transgenic zebrafish line Tg(hsp70l:id3-2A-NLS-KikGR), which carries a heat shock inducible construct that overexpresses id3 after a temperature shift (Row et al., 2018). We previously showed that overexpression of id3 from this line can cause transplanted cells from a donor embryo to faithfully adopt an endothelial fate in a wild-type host embryo. (Row et al., 2018). We utilized this by crossing Tg(hsp70l:id3-2A-NLS- KikGR)), hereafter referred to HS:id3, to HS:dnRAR, to generate donor embryos wherein transplanted cells would be targeted to the endothelium and express dominant negative retinoic acid receptors (RARA). Cells positive for both transgenic constructs were transplanted into tg(kdrl:eGFP) host embryos and these cells exhibited normal migration to the midline (Figure 3A, Supplemental Movie 4). In addition, we utilized a RA reporter line, tg(RDBD,5XUAS:GFP), to determine which tissues were subject to RA signaling (Mandal et al., 2013). We performed immunohistochemistry against GFP and the transcription factor Etv2, which labels the angioblasts (Figure 3B) (Sumanas et al., 2005). The GFP expression indicates tissues that have been exposed to RA. These tissues included the adaxial region of the somites, the notochord, and epidermis (Figure 3B, white arrow, white arrowhead, and gold arrow, respectively). The primitive angioblasts, labeled with anti-Etv2 staining, do not overlap with GFP staining (Figure 3B, red arrows). Together these results indicate that angioblasts are not receiving an RA signal and do not require RA signaling for migration when surrounded by a wild-type environment.

We investigated which tissue was responsible for the angioblast migration defect based on the observed activity of the reporter. Given its proximity to the midline, we chose to test whether the notochord or somitic mesoderm contributed to the defect. To test notochord contribution, we transplanted cells from a wild-type donor to a tg(kdrl:eGFP) host embryo to target the midline progenitors (Row et al., 2016). Host embryos with wild-type cells transplanted to the notochord displayed normal angioblast migration to the midline (Figure 2E). Similarly, transplants of tg(HS:dnRAR) cells into the notochord showed normal midline angioblast migration(Figure 2F). In addition, in-situ hybridizations against a known notochord-secreted angioblast chemoattractant apela, as well as the receptors aplnra and aplnrb, show little change in RA depleted conditions (Figure Supplement 2A-2F). This indicates that the lack of RA signaling in notochord cells is not causing the angioblast migration defects. We then transplanted cells to target the somitic mesoderm. Wild-type cells transplanted into the somitic mesoderm showed normal angioblast migration to the midline (Figure 2G). However, when we transplanted tg(HS:dnRAR) cells into the somitic mesoderm, angioblasts showed migratory defects in the region of the host embryo in which they were transplanted (Figure 2H).

**Figure 2.**
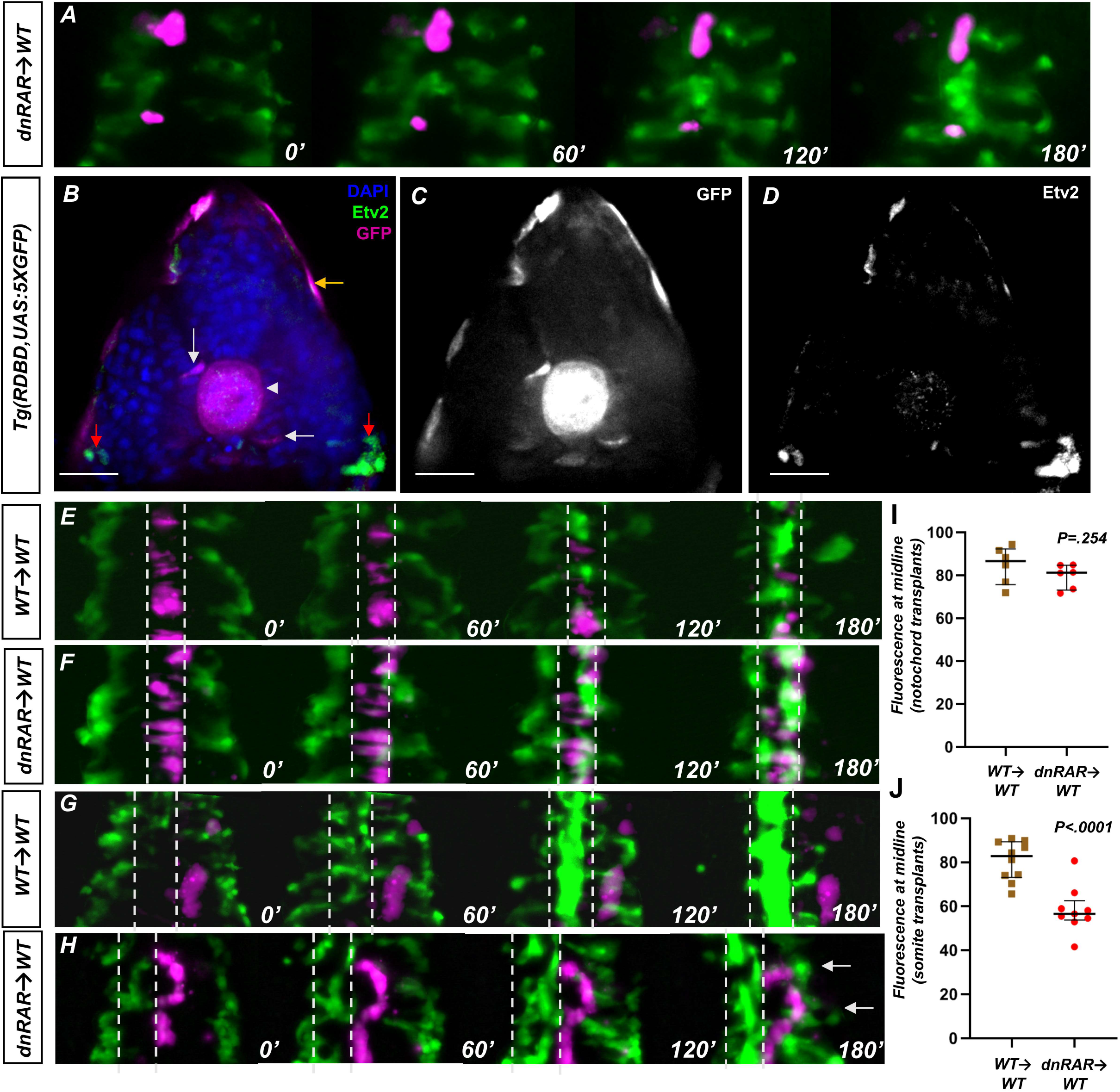
Retinoic acid signaling is not present in the endothelium and is required in the somites for angioblast migration. (A) Representative images of a time-lapse of tg(HS:dnRAR), tg(HS:id3) cells transplanted into a wild-type embryo. Cells labeled in magenta indicate migrating cells. These cells migrate along with the host angioblasts labeled by tg(kdrl:gfp), indicating normal migration patterns in absence of cell-autonomous RA depletion conditions (n=3) (see Supplemental Movie 4). (B) DAPI stained section of 13- somite-stage tg(RDBD,5XUAS:GFP) embryo, immunostained for GFP and Etv2 (n=12). GFP signal is present in adaxial region of the somite, the notochord, and the epidermis (white arrow, white arrowhead, and gold arrows, respectively). The red arrows indicate areas of Etv2 staining. (C) GFP channel of image in (B). (D) Etv2 channel of image in (B). Note the lack of overlap between Etv2 staining and GFP staining. (E) Representative images of a time-lapse of wild-type cells transplanted into a wild-type notochord. (F) Representative images of a time-lapse of tg(HS:dnRAR) cells transplanted into a wild-type notochord. (G) Representative images of a time-lapse of wild-type cells transplanted into a wild-type somite. (H) Representative images of a time-lapse of tg(HS:dnRAR) cells transplanted into a wild-type somite. White arrows indicate angioblast migration defects. (I) Quantification of midline fluorescence, as a percentage of total, for notochord transplants at 180’. The difference for wild-type and tg(HS:dnRAR) were not statistically significant (P=0.254). (J) Quantification of midline fluorescence, as a percentage of total, for somite transplants at 180’. The difference for wild-type and tg(HS:dnRAR) were statistically significant (P<0.0001). Scale bars, 50 µm.

Quantification of fluorescence percentage at the midline, in the chimeric regions of the notochord, showed no statistically significant difference compared to controls (Figure 2I).

However, quantification of fluorescence percentage in the chimeric regions of the somites showed statistical significance between wild-type and tg(HS:dnRAR) (Figure 2J). This indicates that the somites are the principal tissue required for RA signaling mediated midline angioblast migration.

### The somitic mesoderm is required for angioblast migration to the midline

To follow up the somite and notochord targeted transplant results, we examined midline angioblast migration in embryos where these tissues are absent. Previous studies have investigated genes linking the notochord to angioblast migration. Mutants of the genes noto and apela were found to be notochord specific genes required for proper angioblast migration (Cleaver and Krieg, 1998; Fouquet et al., 1997; Helker et al., 2015). The noto gene induces notochord, and the notochord in turn secretes Apela which attracts the angioblasts to the midline. Noto is required for the formation of the notochord, and in the absence of Noto, the midline cells adopt a somitic fate (Talbot et al., 1995).

However, it is not clear if midline convergence was delayed or absent in notochord-less embryos. In noto mutants, some angioblasts reach the midline at later developmental stages, and resolve into a single blood vessel (Fouquet et al., 1997). We injected a noto morpholino that faithfully phenocopies the noto mutant into tg(kdrl:GFP) embryos (Ouyang et al., 2009). Control morphant embryos show normal angioblast migration (Figure 3A), whereas noto morphants show slowed angioblast migration (Figure 3B). However, midline convergence still occurs in both conditions (Figure 3C). This implies that the notochord is ultimately dispensable for the formation of the VMC and midline positioning of angioblasts.

**Figure 3.**
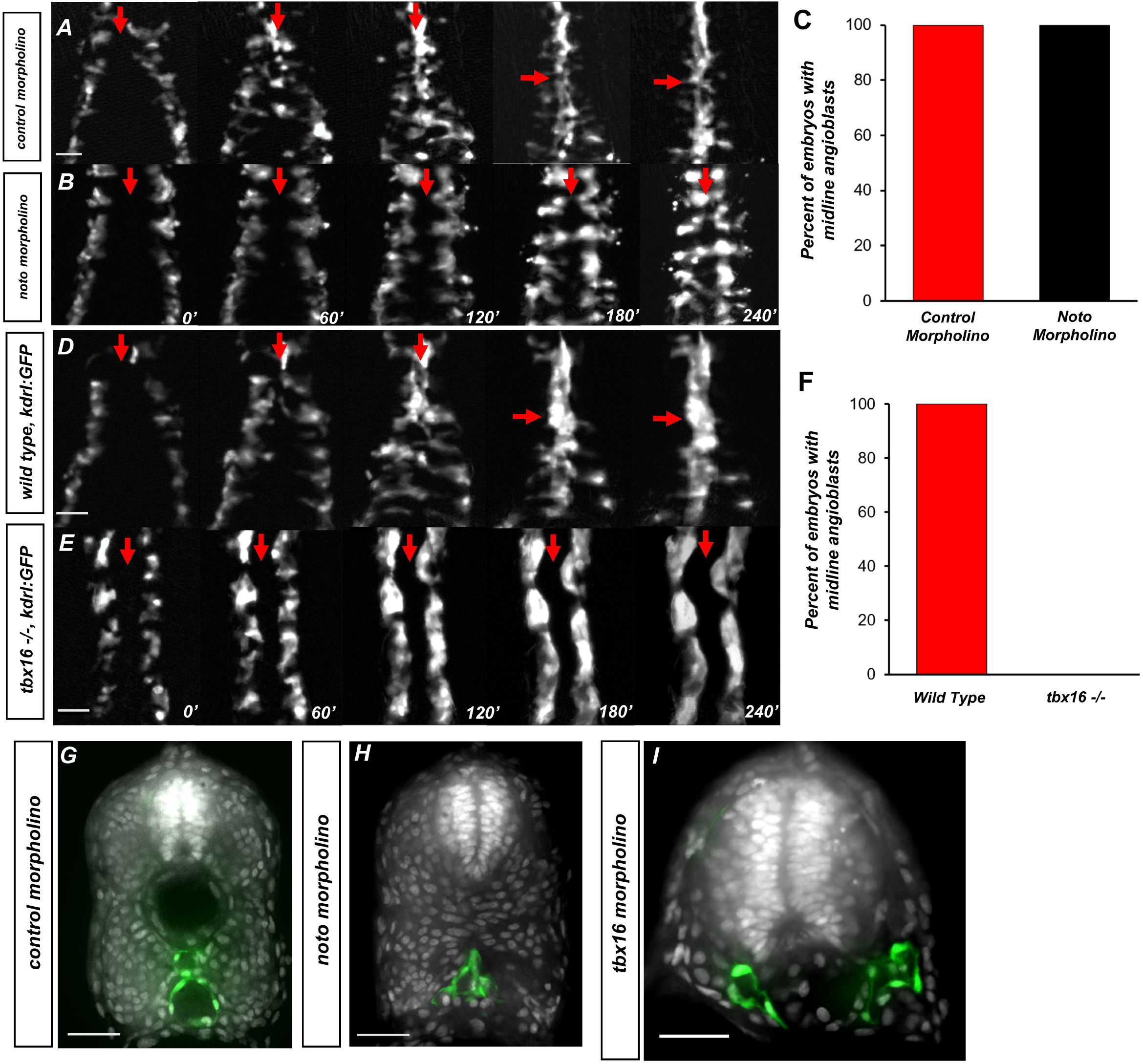
The somitic mesoderm, not the notochord, is required for midline convergence of angioblasts. (A) Time-lapse imaging of tg(kdrl:eGFP) embryos injected with control morpholino over a 240’ period. (B) tg(kdrl:eGFP) embryos injected with noto morpholino over a 240’ period. Red arrows indicate the midline. (C) Graph of embryos with midline angioblasts in noto and control morphants after 240’. N=15 for both conditions (D) Time-lapse imaging of wild-type, tg(kdrl:eGFP) embryos over a 240’ period. (E) Timelapse imaging of tbx16 -/-, tg(kdrl:eGFP) embryos over a 240’ period. Red arrows indicate the midline. (F) Graph of embryos with midline angioblasts in wildtype and tbx16 -/- after 240’ N =18 for both conditions. (G) Section of wild-type, tg(kdrl:eGFP) embryos over a 240’ period. (H) Section of 24 hpf tg(kdrl:eGFP) embryos injected noto morpholino and stained with DAPI. (I) Section of 24 hpf tg(kdrl:eGFP) embryos injected tbx16 morpholino. Blood vessels are labeled in green and nuclei labeled in grey. Scale bars, 50 μm.

Having ruled out the role of notochord tissue, we next investigated whether the somitic mesoderm is required for NES and the formation of the VMC. Previous studies have shown that the t-box transcription factor tbx16 is required for proper somite formation (Amacher et al., 2002; Goto et al., 2017; Griffin et al., 1998; Kimmel et al., 1989; Manning and Kimelman, 2015; Row et al., 2011). In the absence of Tbx16, cells that would normally join the paraxial mesoderm fail to do so, causing a deficiency in the trunk somitic mesoderm, but leaving the endothelium intact (Thompson et al., 1998). We utilized tbx16 mutants to determine whether the lack of somites caused angioblast migration defects, similar to RA loss of function. We examined midline angioblast migration in tbx16 mutant embryos with the tg(kdrl:GFP) transgene in the background. In wild-type sibling embryos, angioblast migration progresses normally to the midline (Figure 3D). However, in tbx16 -/- embryos, angioblast migration does not occur, and they remain in their positions in the lateral plate (Figure 6E). Fluorescence quantification indicated that angioblasts never reach the midline in any embryo (Figure 3F). To determine the effect of either notochord or somite loss on the definitive vasculature, we performed transverse sections of tg(kdrl:GFP) embryos injected with control, noto, and tbx16 morpholinos and stained with DAPI. Control morpholino showed normal formation of the dorsal aorta and common cardinal vein (Figure 3G). Embryos injected with noto morpholino have angioblasts located at the midline that appear to have resolved into one blood vessel structure (Figure 3H). Embryos injected with tbx16 morpholinos, however, show two distinct vessels located on either side of the notochord (Figure 3I). This indicates that the somites are critical for angioblast convergence at the midline, while the notochord is ultimately dispensable.

### Retinoic acid induces a dorsal translocation of the notochord away from the underlying endoderm

Given the non-autonomous midline angioblast migration defects in RA loss of function embryos, we examined if RA manipulation altered the normal development of non- angioblast midline tissues. We utilized embryos labeled with a cell surface marker, tg(ubb:lck-mng) (Adikes et al., 2020), and found the relative location of notochord at the midline changes based on retinoic acid activity. To measure this, embryos were treated with either DMSO, DEAB, or RA at the tailbud stage. The embryos were then fixed, deyolked, and imaged in the trunk at roughly the 5^th^ somite (Figure 4A-C). The relative location of the notochord in DMSO treated embryos was in close proximity to the ventral underlying endoderm at the 12-somite stage, but that distance increased by the 15- somite stage (Figure 4A). In DEAB treated embryos, the notochord remains ventrally localized at both the 12-somite stage and 15-somite stage (Figure 4B). When the same experiment was done with embryos exposed to exogenous RA, the notochord was prematurely dorsally localized at the 12-somite stage and extended even further by the 15-somite stage (Figure 4C).

**Figure 4:**
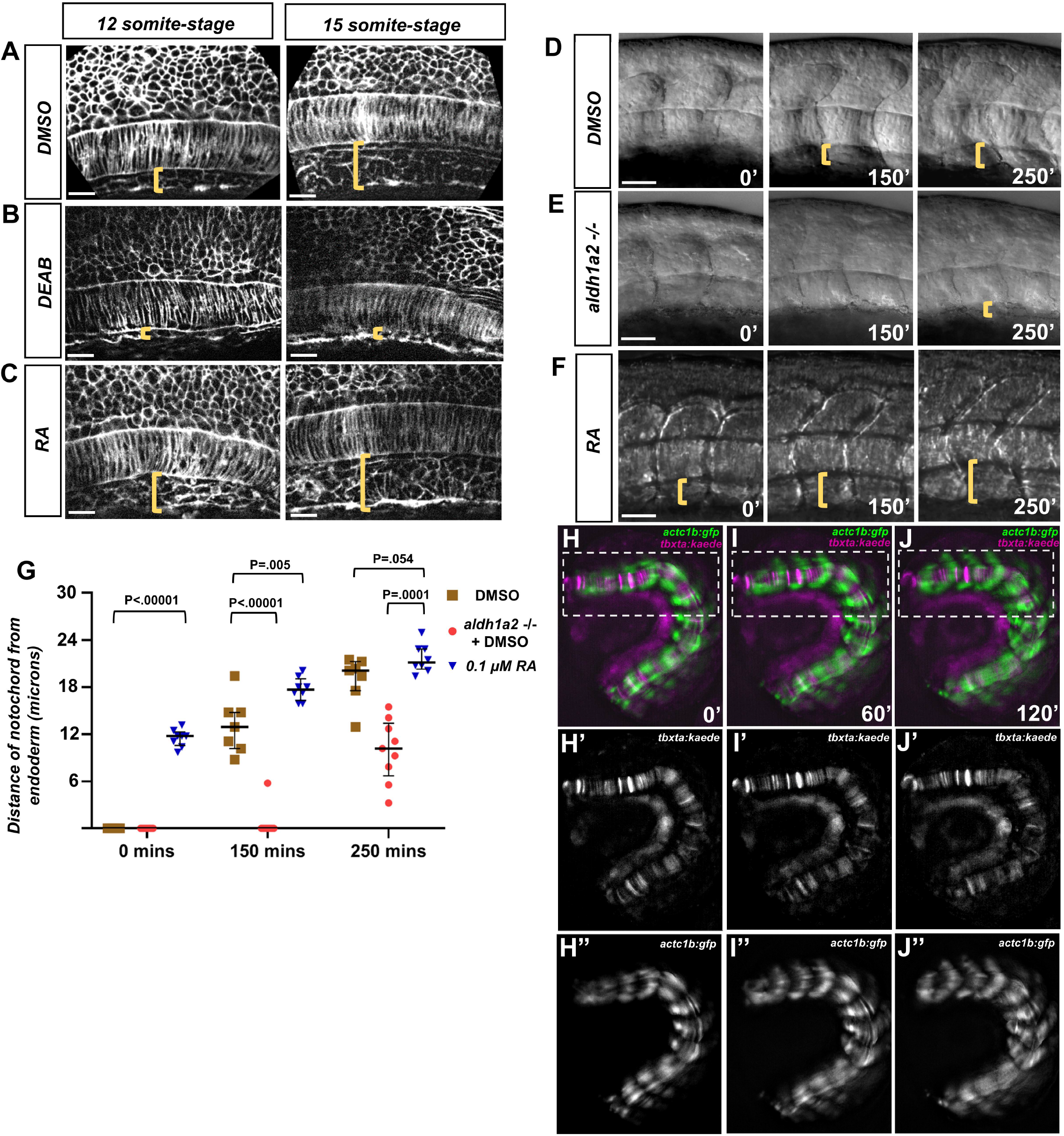
RA signaling promotes dorsal translocation of the notochord. (A-C) Fluorescent images of fixed (A) DMSO treated (B) DEAD treated and (C) RA treated tg(ubb:lck-mng) embryo trunks at the 12- and 15-somite stage. Yellow brackets indicate the distance between the notochord and underlying yolk cell. (D-F) DIC time-lapse images of the notochord and zebrafish trunk. Focal planes of the notochord and somites were overlaid to show the relative location to one another. Yellow brackets indicate the distance of the ventral notochord to the ventral-most portion of the embryo in (D) DMSO treated embryos, (E) aldh1a2 -/- embryos treated with DMSO, or (F) 0.1 μM RA treated embryos. (G) Quantification of dorsal translocation of notochord from the ventral tissue. Black lines indicate the median and interquartile range. Wild- type are indicated by brown squares, while aldh1a2 -/- and 0.1 μM RA embryos are shown by red circles and blue triangles, respectively. At t=0, DMSO vs. RA is P<.00001. At 150’, DMSO is P<.00001 and P=.005 vs. aldh1a2 -/- and RA treatment, respectively. At 250’, DMSO is P=.0001 and .054 vs. aldh1a2 -/- and RA treatment, respectively. (H-J) Time-lapse image of a trunk explant from tg(actc1b:gfp), tg(tbxta:kaede) embryo at (H) 0’, (I) 60’, and (J) 120’. (H’-J’) Time-lapse image of explant showing tg(tbxta:kaede) only. (H’’-J’’) Time-lapse image of explant showing tg(actc1b:gfp) only. Time-lapse shows the notochord adjacent the yolk at time 0’, but moves dorsally from the yolk at time 120’. Scale bars, 25 µm.

We then determined the dynamics of the notochord displacement in real time. Using DIC images, we were able to overlay the lateral trunk with the notochord. We then observed the notochord displacement in a 250’ time-lapse for DMSO treated wild-type, aldh1a2 -/- with DMSO, and exogenous RA (Figure 4D, 4E, and 4F, respectively). We quantified this data by measuring the distance of the ventral aspect of the notochord to the yolk cells of the embryo (yellow brackets). We found that aldh1a2 -/- embryos showed delayed dorsal displacement relative to DMSO in both the timing of dorsal movement and the magnitude of that movement (Figure 4G).

Previous work showed that as somites mature, they extend in the both the dorsal and ventral axis (Tlili et al., 2019), and based on this we hypothesized this extension was concurrent with the dorsal translocation of the notochord. However, we wanted to rule out the effect of forces on the anterior-most and posterior-most regions of the notochord. The notochord is a rigid structure that is anchored in the tailbud near the notochord primordia, and extends into the head mesoderm medial to the otic vesicle (Kimmel et al., 1995). To eliminate the contribution other tissues could have on NES, we generated trunk explants from embryos containing both tg(actc1b:gfp) and tg(tbxta:kaede) transgenese that contained roughly 10 somites of the embryo. The 10-somite stage embryos were sectioned near the somite borders of the 1^st^ and 10^th^ somite. We performed live imaging of these trunk explants, focusing on the immature somites of the posterior region (white boxes) (Figure 4H-4J, Supplemental Movie 5). In order to better visualize the notochord, we separated the tg(actc1b:gfp) and tg(tbxta:kaede) signals into separate channels. This distinguished the notochord reporter tg(tbxta:kaede) (Figure 4H’-4J’), from the somite reporter tg(actc1b:gfp) (Figure 4H”-4J”). The notochord reporter showed clear dorsal movement away from the auto-fluorescent yolk (Figure 4H’-J’). The somites also showed the corresponding expansion along the dorsal-ventral axis (Figure 4H”-4J”). Together, this data indicates that the proper development of the midline is both retinoic acid-dependent and requires only factors localized to the trunk of the embryo.

### Notochord-endoderm separation facilitates the midline convergence of angioblasts

We speculated that RA dependent changes at the midline could be the cause of the angioblast migration defect. In order to test this, we generated a heat-shock inducible transgenic reporter of F-actin, tg(hsp70l:lifeact-mScarlet), which labels filamentous actin throughout the embryo and allows visualization of all cells (Riedl et al., 2008). We hereafter refer to tg(hsp70l:lifeact-mScarlet) as tg(HS:lifeact). We then crossed this reporter to the tg(kdrl:eGFP) line to label the angioblasts. We treated embryos with either DMSO (Figure 5A-5C), exogenous RA (Figure 5D and 5E), or DEAB (Figure 5F- 5I). Embryos were sectioned at the region of the 5^th^ somite (Figure 5A-5I). Embryos that were treated with DMSO showed normal midline angioblast migration patterns from the 12-somite stage, 15-somite stage, and 18-somite stage (Figure 5A-5C). As angioblasts approach the midline, a gap appears between notochord/hypochord and the underlying endoderm (Figure 5A). This gap is filled by angioblasts by the 15-somite stage, separating the notochord from the endoderm (Figure 5B). We call the process of dorsal translocation of the notochord the notochord-endoderm separation (NES). The transient opening that forms from NES is referred to as the ventral midline cavity (VMC). As angioblast migration continues, additional angioblasts occupy the VMC until the 18- somite stage, when angioblast migration is largely complete for the 5^th^ somite region of the trunk (Figure 5C).

**Figure 5.**
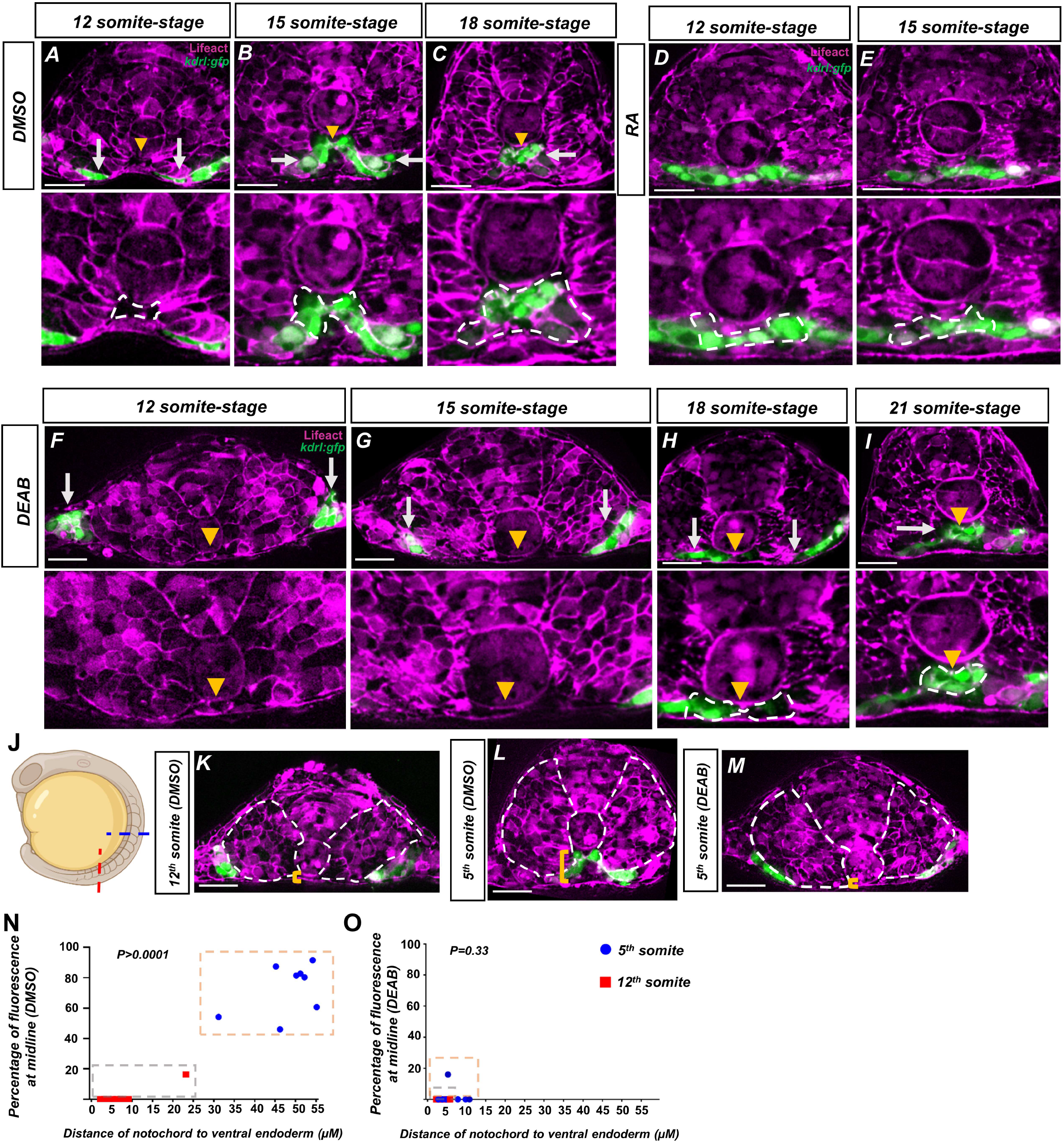
Retinoic acid mediates a notochord-endoderm separation required for terminal angioblast migration. (A-I) Embryos generated by crossing tg(HS:lifeact) to tg(kdrl:eGFP) to label both actin and angioblasts. Yellow arrowheads indicate the midline while white arrows indicate angioblasts. A magnified image includes a white dashed line to indicate notochord-endoderm separation. (A-C)) DMSO treated embryos sectioned at the 5^th^ somite during the (A) 12-somite stage (n=8), (B) 15- somite stage (N=8), and (C) 18-somite stage (N=6). (D-E) 0.1 μM RA treated embryos sectioned at the 5^th^ somite during the (D) 12-somite stage (N=8) and (E) 15-somite stage (n=8). (F-I) 20 μM DEAB treated embryos sectioned at the 5^th^ somite during the (F) 12-somite stage (N=7), (G) the 15-somite stage (N=8), (H)18-somite stage (N=6), and (I) 21-somite stage (N=6). (J-L) 15 somite- stage embryos generated by crossing tg(HS:lifeact) to tg(Kdrl:eGFP). White dashed lines indicate outline of somites while yellow brackets indicate distance of notochord to ventral endoderm. (J) Schematic showing the sectioned region for experiments in (K-O). Red line indicates sectioning at 5^th^ somite and the blue line indicates sectioning at the 12^th^ somite. (K) A DMSO embryo sectioned at the 12^th^ somite. Note the lack of midline angioblasts and small distance between the notochord and the ventral endoderm. (L) A DMSO-treated embryo sectioned at the 5^th^ somite. Note the angioblasts at the midline and larger distance from the notochord to the ventral endoderm. (M) A DEAB embryo sectioned at the 5^th^ somite. Note the lack of midline angioblasts and small distance between the notochord and the ventral endoderm. (N) A two variable graph showing percentage of angioblast fluorescence at the midline compared to distance between the notochord and the ventral endoderm. Blue dots indicate DMSO treated, 15-somite stage embryos sectioned at the 5^th^ somite and red squares are the same embryos sectioned at the 12^th^ somite (N=8). (O) A two variable graph showing percentage of angioblast fluorescence at the midline compared to distance between the notochord and the ventral endoderm. Blue dots indicate DEAB treated, 15-somite stage embryos sectioned at the 5^th^ somite and red squares are the same embryos sectioned at the 12^th^ somite (N=8). Scale bars, 50 µm.

To determine if activation of retinoic acid is sufficient to accelerate angioblast migration and NES concurrently, we treated embryos with exogenous RA and sectioned them at the 12-somite and 15-somite stage. Embryos sectioned at the 12-somite stage showed premature angioblast migration to the midline and NES, indicating that angioblast migration and NES are responsive to exogenous RA signal (Figure 5D). Angioblasts continue to be at the midline at 15-somite stage (Figure 5E). 18-somite stage embryos were not available as posterior elongation discontinued in prolonged RA exposure, as previously shown (Martin and Kimelman, 2010). To determine whether angioblast migration and NES was attenuated in RA depletion conditions embryos were treated with DEAB and sectioned at the 12-somite, 15-somite, 18-somite, and 21-somite stage (Figure 5F-5I respectively). At the 12-somite stage, the angioblasts are localized in the lateral plate mesoderm (Figure 5F). Focusing on the midline of the embryo, we observe that the notochord remains flush against the endoderm with no visible VMC or NES (Figure 5G). At the 15-somite stage, we observe some midline movement of the angioblasts, however the notochord remains flush against the endoderm and no angioblasts have been able to reach the midline (Figure 5G). During the 18-somite stage, a small VMC has formed between the notochord and endoderm. The leading angioblasts have partially migrated into this VMC, however the bulk of the angioblasts still reside outside of the midline (Figure 5H). By the 21-somite stage, the angioblasts have largely resolved to the midline, although with some angioblasts still residing outside the midline (Figure 5I).

Given the pronounced delays in NES formation for anterior regions in RA depleted embryos, we suspected that NES was a consequence of the known processive somite maturation that begins in the anterior regions and progresses posteriorly (Stern and Piatkowska, 2015). We sectioned 15-somite stage tg(HS:lifeact), tg(kdrl:egfp) embryos in the anterior and posterior regions corresponding to the 5^th^ and 12^th^ somite (Figure 5J). In more immature 12^th^ somite, angioblasts reside in their lateral regions and NES is not pronounced (white dashed lines outline the somites, and yellow brackets show the distance from the notochord to the ventral endoderm) (Figure 5K). In the 5^th^ somite of the same embryo, angioblasts reside at the midline and NES is very pronounced, indicating NES occurs anterior to posteriorly (Figure 5L). The lack of angioblasts at the 12^th^ somite, and low NES, is similar to DEAB treated embryos sectioned at the more mature 5^th^ somite position (Figure 5M).

To quantify how the dorsal translocation of the notochord was correlated with angioblasts arriving to the midline, We created a two variable graph to measure both the distance of the notochord to the ventral endoderm and the percentage of angioblasts arriving at the midline for a given somite. For this analysis, we used 15-somite stage embryos sectioned at the 5^th^ somite and 12^th^ somite. We expect that immature somites in the posterior would have little midline migration of angioblasts and the anterior, with more mature somites, would have significant midline migration. We measured the distance in microns on the x-axis and the percentage of midline angioblasts on the y-axis (Figure 5O-5P). In DMSO treated embryos, embryos sectioned at the 5^th^ somite grouped together, showing both significant notochord-endoderm distance and angioblasts residing at the midline (Figure 5O, blue dots in the orange box). Embryos sectioned at the 12^th^ somite show short notochord-endoderm distance and little angioblast migration towards the midline (Figure 5O, red squares in grey box). When comparing angioblasts migration, the 5^th^ somite and 12^th^ somite regions were significantly different (P>.0001).

The corresponding sectioning of DEAB treated embryos shows a difference compared to wild-type in the 5^th^ somite region. Embryos sectioned at the 5^th^ somite have a small distance from the notochord to endoderm, as well as no angioblast migration to the midline (Figure 5P, blue dots in orange box). Interestingly, the notochord-endoderm distance/migration percentage ratio strongly resembled the more immature somites. The angioblast migration percentage was not statistically significant between the 5^th^ and 12^th^ somite in RA depletion conditions (P=.33). This indicates that the midline exists in a more immature state in RA depleted conditions, and the more mature state provides space in which the angioblasts can migrate into.

### Retinoic acid induces changes in the definitive vasculature

Pharmacological inhibition of RA signaling has previously been shown to effect the zebrafish vasculature, notably causing a smaller than normal dorsal aorta (Pillay et al., 2016). To confirm this effect in the aldh1a2 -/- mutant, we sectioned 24 hpf tg(kdrl:GFP) and aldh1a2 -/- embryos, stained the sections with DAPI, and imaged them using spinning disk confocal microscopy (Figure 6A, 6B). Images show that the dorsal aorta and posterior cardinal vein in wild-type embryos are roughly equivalent in size (Figure 6A, white arrows). However, in the aldh1a2 -/- embryos, the size of the dorsal aorta is reduced but remained lumenized (Figure 6B, white arrows). We confirmed this observation by in-situ hybridization of the arterial marker cldn5b and the venous marker dab2, which showed similar effects (Figure S2A-D) (Casie Chetty et al., 2017). To determine if this trend extended beyond the 24 hpf developmental stage, we imaged the trunk of 5 dpf tg(kdrl:GFP) embryos along with tg(kdrl:GFP), aldh1a2 -/- siblings (Figure 6C,6D). The tg(kdrl:GFP) embryos alone showed no vascular defects and normal dorsal aorta and posterior cardinal vein (Figure 6C, white arrow and yellow arrowhead respectively). However, the vasculature of the aldh1a2 -/- siblings show a large reduction of the size of the dorsal aorta (Figure 6D, white arrow). In addition, it appears that the posterior cardinal vein of aldh1a2 -/-increased in size compared to its sibling (Figure 6D, yellow arrowhead). To determine if the size change was the result of a reduced cell number in the DA in response to RA loss, we utilized the transgenic line tg(kdrl:nls-GFP), which labels the nucleus of endothelial cells to count the number of cells in each blood vessel over a 225 μM horizontal section of a 24 hpf zebrafish embryo. Embryos were treated with either DMSO, or with DEAB to inhibit RA. We see reduced cell number in the DEAB treated artery versus the DMSO treated artery (Figure 6E). This corresponds to an increase number of cells in the veins during DEAB treatment compared to DMSO. While we cannot definitively say the migration defects caused the arterial reduction, it indicates that RA is required for the proper formation of the definitive vasculature.

**Figure 6.**
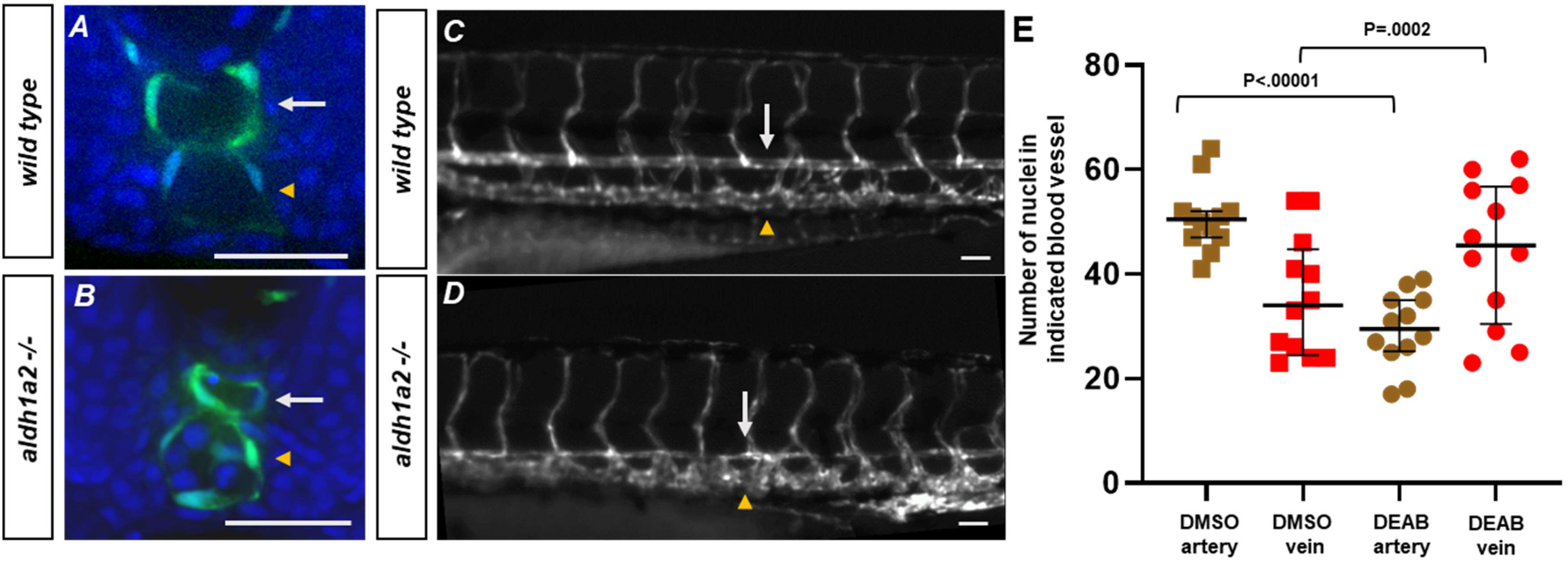
Retinoic acid loss contributes to changes in the definitive vasculature, resulting in large veins and small arteries. (A) A 5 dpf tg(kdrl:eGFP) embryo. White arrows indicate the dorsal aorta and yellow arrowheads indicate the cardinal vein. (B) 5 dpf tg(kdrl:eGFP), aldh1a2-/- embryo. White arrows indicate reduced size of dorsal aorta. (C-D) 25 hpf tg(kdrl:eGFP). embryo labeled with DAPI. (C) Wild-type embryos show lumenized blood vessels and normal sized dorsal aorta. (D) Labeled aldh1a2 -/-, tg(kdrl:eGFP) sibling shows small lumenized dorsal aorta with large posterior cardinal vein. (E) Quantification of DMSO (n=12) or 20 μM DEAB (n=12) treated tg(kdrl:nls-eGFP) embryos. Nuclei were counted in each structure over a 225 μM long transverse section of the zebrafish trunk. Two-tailed P-values for the unpaired t-test of wild-type vs 20 μM DEAB treated embryos are <.00001 and .0002 for arteries and veins, respectively. Scale bars, 50 μm.

### Retinoic acid induced somite morphogenesis facilitates notochord-endoderm separation and facilitates midline angioblast migration

While NES and midline formation were shown to be highly correlated, we previously showed that cell autonomous RA depletion within the somite caused angioblast migration defects. Therefore, we sought to determine if RA depletion within the somites caused morphological changes that compromised NES and angioblast migration to the midline. During normal development, newly-born somites initially lay flat and extend farther in the medial-lateral axis, demonstrated by a section of 15-somite stage embryo at the 12^th^ somite (Figure 7A, white dashed lines) (Tlili et al., 2019). However, as somites mature, they will narrow in the medial-lateral axis and extend in the dorsal-ventral axis, as demonstrated by a section of the 5^th^ somite of a 15-somite stage embryo (Figure 7B, white dashed line) (Tlili et al., 2019). We therefore sought to determine if somite maturation could be affected cell autonomously in RA depletion conditions. We therefore transplanted either wild-type or HS:dnRAR cells into tg(ubb:lck-mng) embryos. Embryos were then sectioned at roughly the 5^th^ somite region for 12-somite stage embryos (Figure 7C and 7D). Embryos that were transplanted with wild-type cells showed little difference with the contralateral somite in terms of shape and maturation (Figure 7C, white dashed lines). However, embryos transplanted with HS:dnRAR cells showed a contralateral defect in the somite containing donor cells, with the somite extending farther in the medial-lateral axis while the contralateral somite extends farther in the dorsal-ventral axis (Figure 7D, white dashed lines). To illustrate this effect across multiple experimental embryos, we show overlapping shapes of chimeric wild-type somites, the chimeric HS:dnRAR somites, and the contralateral somites from HS:dnRAR chimeras (Figure 7E). The overlapping images show a consistent trend of a shallow dorsal-ventral axis in the chimeric HS:dnRAR somites compared to the wild-type somites. Together, this indicates that RA is required cell autonomously for somite maturation and dorsal-ventral extension.

**Figure 7.**
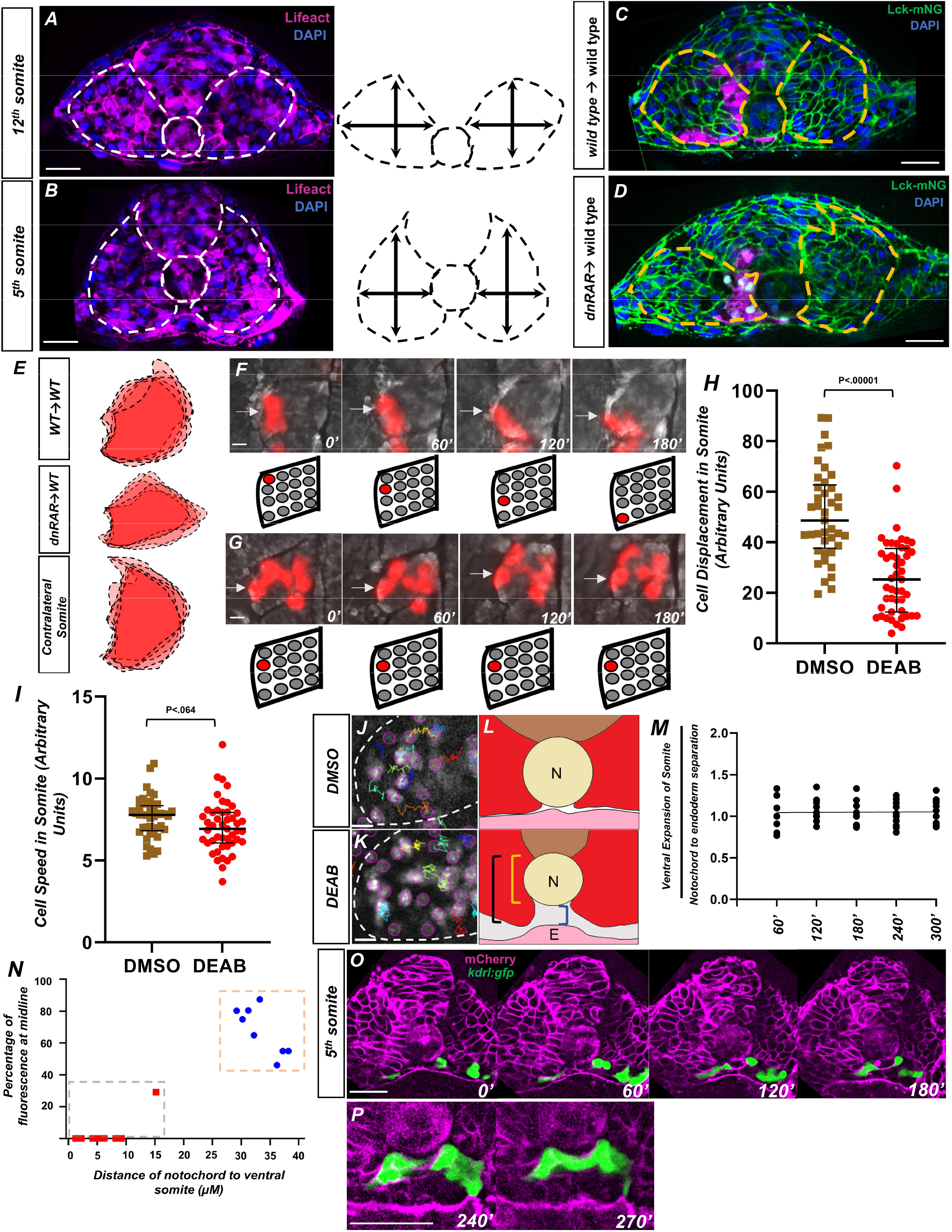
Retinoic acid controls intrasomitic cellular movements and somite shape changes that contribute to angioblast convergence to the midline. (A) Image and schematic of newly born 12^th^ somite using tg(HS:lifeact), tg(kdrl:egfp) embryo as reference. Dashed lines indicate the somite and notochord. (B) Schematic of more mature 5^th^ somite (at the 12-somite stage) of tg(HS:lifeact), tg(kdrl:egfp) embryo as reference. Dashed lines indicate somite and notochord. (C) Transplant of wild-type cells into tg(ubb:lck-mng). Dashed lines indicate somite shape. (D) Transplant of tg(HS:dnrar) cells into tg(ubb:lck-mng). Dashed lines indicate somite shape. Note the immature shape of the somite containing transplanted cells. (E) Overlapping somite shapes of either WT→WT, tg(HS:dnrar)→WT, or the contralateral somite tg(HS:dnrar)→WT. (F-G) Time-lapse images of transplanted rhodamine-dextran labeled cells in DMSO and DEAB embryos at 12 somite stage. (F) DMSO treated embryos over 180’. White arrows indicate labeled cells (G) 20 μM DEAB treated embryos over 180’. Scale bars, 50 μm. Retinoic acid depleted embryos show little cellular movement during somite development. (H) Quantification of nuclei displacement in the 2^nd^ somite of DMSO and DEAB embryos over a 4 hour time period. Displacement is defined as the distance between the beginning location and end location of the nuclei within the somite 2. (I) Quantification of speed in somite 2, defined as average units over 4 hours in 10 minute increments. (J) Sample of tracks from DMSO treated tg(hsp70l:CAAX-mCherry-2A-NLS-KikGR) embryos. (K) Sample tracks of DEAB treated tg(hsp70l:CAAX-mCherry-2A-NLS-KikGR) embryos. Tracks sort from red to blue, with red being the highest displacement. Scale bars, 20 µm (L) Diagram of ventral somite expansion and notochord to endoderm separation. Ventral somite expansion is calculated by ventral somite length (black bracket) – the notochord diameter (Yellow Bracket). Notochord/endoderm separation is shown in the blue bracket. (M) Quantification of the ventral expansion of the somite over notochord/endoderm separation in wild-type embryos. The linear regression indicated by the black line, where slope = 0.002, shows ventral somite expansion correlates with notochord/endoderm separation. (N) A two variable graph showing percentage of angioblast fluorescence at the midline compared to distance of ventral somite expansion beneath the notochord. Blue dots indicate DEAB treated, 15-somite stage embryos sectioned at the 5th somite and red squares are the same embryos sectioned at the 12th somite (N=8). Scale bars, 50 µm. (O) Live section of mcherry-caax injected tg(kdrl:egfp) embryos over 120’. Yellow arrowheads indicate the midline while white arrows indicate angioblasts. (P) Same live section as in (N), except at 240’ focused at the midline. Note the separation of the hypochord and endoderm prior to midline fusion of angioblasts. Scale bars, 50 µm.

We speculate that the failure of somites to mature was the result of reduced cell movements within the somite. To confirm this, mosaically labeled cells were generated by transplanting rhodamine dextran labeled cells from a wild-type donor into a wild-type host. The embryos were treated with either DMSO or DEAB and time-lapse imaged over the course of 4 hours (Figure 7F and Figure 7G, respectively). Cells in DMSO treated embryos showed broad movements consistent with that of somite rotation, where cells located in the anterior of the somite will translocate to the posterior of the somite (Hollway et al., 2007) (Figure 7F, white arrows). However, cells in DEAB treated embryos show little intratissue displacement and cells largely remained static (Figure 7G, white arrows). To quantify this, we utilized tg(hsp70l:CAAX-mCherry-2A-NLS-KikGR) embryos to observe cell movement across the entirety of the somite. Embryos were heat shocked during shield stage and subsequently treated with either DMSO or DEAB and their somites were time-lapse imaged for 4 hours. Using Fiji Trackmate software (Tinevez et al., 2017), we quantified the displacement and speed of cell movements within the 2nd somite (Figure 7H and 7I). Cells in DMSO treated embryos showed significant amounts of displacement throughout the somite (Figure 7H).

However the DEAB cells showed less displacement over time (Figure 7H). Quantification of speed in the same manner showed no statistically significant difference between DMSO and DEAB treatment (Figure 7I). A sample of somite cell tracks shows little directional movement in loss of RA conditions compared to DMSO, indicating cells are not moving in a manner consistent with reshaping of the somite, as well as suggesting a role for RA signaling in somite rotation (Figure 7J and 7K, Supplemental Movie 6 and 7, respectively). The Cxcr4a/Cxcl12a signaling axis has been shown to be responsible for controlling cell movements in the somite. In-situ hybridization for cxcr4a shows a loss of expression in RA depletion conditions and gain of expression in RA addition conditions (Hollway et al., 2007) (Figure S5D and S5E).

We speculated that the ventral expansion of the somite that occurs during somite cell rearrangement and maturation was responsible for NES. At the 10 somite stage, embryos have their notochord adjacent to the endoderm and level to ventral-most edge of the somite (Figure 7L). As the embryo develops, the somite expands ventrally (black bracket). As the ventral distance of the somite extends relative to the dorsal part of the notochord (yellow bracket), it induces space that the angioblasts occupy (blue brackets). Starting from the 10 somite stage, we measured the distance of ventral expansion relative to the distance between the notochord and the endoderm every hour for 4 hours. A simple linear regression of the data, indicated by the black line, shows a slope of 0.2 and y-intercept of 1.04 (Figure 7M). The flat slope indicates little variance across time, and the y-intercept near 1 indicates that NES is equivalent to ventral expansion of the somite. The 1:1 correlation of ventral somite expansion and notochord displacement, as well as the tbx16 loss of function NES defect, provides strong evidence that somite morphogenesis is critical for NES.

Finally, we investigated a methodology to track all cellular movements within an embryo to co-visualize somite morphogenesis along with angioblast migration and notochord- endoderm separation. We used a live explant method to monitor migration in a transverse section view. Live tg(kdrl:GFP) embryos, injected with mcherry-caax mRNA, were sectioned at the 10-somite stage at the 4^th^ somite and mounted to observe a transverse section of the trunk. We were able to observe migrating angioblasts, as well as somite morphogenesis and midline NES in the explant (Figure 7O-7P, Supplemental Video 8). At the 180’ time-point, angioblasts were not at the midline and the hypochord remained flush with the endoderm (Figure 7O, Supplemental Video 8). At this time-point the somites have begun to narrow in the medial-lateral axis. By the 240’ stage, the hypochord is separated away from the endoderm (Figure 7P). and shortly after this, the angioblasts are able to complete their migration to the midline by the 270’ time-point (Figure 7P). Similar to the results seen in fixed sections, live explants of embryos treated with DEAB show angioblast migration defects and lack of NES (Supplemental Video 9).

## DISCUSSION

Here, we established a cell nonautonomous angioblast migration model wherein shape changes in the somitic mesoderm create a physical space for angioblasts to migrate into and develop into the axial vasculature. The emergence of the transient VMC is a necessary requirement for contralateral angioblasts to complete their migration and coalesce at the midline. We observed a defect in somitic mesoderm development in RA depletion conditions, which delayed somite maturation, intrasomite cellular movement, and significantly impeded NES. Conversely, we found that addition of RA accelerated somite maturation and NES, along with angioblast migration. We were able to confirm that the somitic mesoderm was required for angioblast migration by examining tbx16 mutants, which have a lack of somites caused by EMT defects (Goto et al., 2017; Ho and Kane, 1990; Manning and Kimelman, 2015; Row et al., 2011). Loss of function of tbx16 prevented angioblast migration completely and resulted in bifurcated blood vessel formation. These changes in vascular morphology indicate that the somitic mesoderm is a requirement for angioblast migration to the midline. It is notable that in amniotes, such as the mouse and quail, the dorsal aorta and cardinal vein are initially bifurcated in a similar manner to zebrafish tbx16 mutants prior to fusion at the midline (Drake and Fleming, 2000; Pardanaud et al., 1996, 1987). It is possible that evolutionary differences in the somitic mesoderm between teleosts and amniotes could play a role in the differences in vascular development.

Very little is known about the process of NES. Our observations indicate that its initiation correlates very closely with angioblast migration, with angioblasts migrating into the VMC seemingly as soon as it appears in both wild-type and RA depletion conditions.

Given that these blood vessels become lumenized, and need to achieve a sufficient volume for blood transport, it is plausible that they require morphological changes within the ventral part of the embryo to accommodate for their size. Delays in somite maturation prevent angioblasts from reaching the midline in a normal developmental time frame. It is possible that the consequences of this migration defect results in the malformed blood vessel structure in RA depleted embryos, which have small arteries and larger veins. Prior work showed that the most lateral angioblasts, which migrate later than more medial angioblasts, preferentially adopt a venous fate over an arterial one (Kohli et al., 2013). This implies that somite maturation is a key factor in determining the proper distribution of endothelial progenitors to the arteries and veins, by essentially timing angioblast migration and altering cell fate decision patterns. However, we cannot definitively determine if this is solely from angioblast migration defects.

We found that the formation of the VMC does not necessarily require the notochord tissue itself. In noto morphants, which causes the notochord to adopt a somitic fate, there is still a ventral cavity and angioblasts are able to arrive at the midline. This implies that the notochord is ultimately dispensable for the completion of angioblast migration, even though its loss can produce migration defects (Helker et al., 2015; Sumoy et al., 1997). The migratory defects could be the result of reduced notochord-secreted factors such as apela, which greatly reduce but do not completely abolish migration (Helker et al., 2015). Another factor could be that signals from the notochord, such as Hedgehog, influence somite development which in turn facilitates angioblast migration (Blagden et al., 1997; Hinits et al., 2009; Yin et al., 2018). While the fate of the midline tissue does not need to be notochord for NES to occur, it is likely that mechanical coupling between the somites and midline tissue is required such that somite morphogenesis drives the separation of the midline mesoderm away from the underlying endoderm. In noto loss of function, the somitic tissue that forms at the midline in place of notochord is continuous with the flanking somites. In wild-type embryos, there is evidence of mechanical interactions of the somite and notochord during convergent extension, which are required for the remodeling of the adaxial cells, the medial most cells in the somite (Yin and Solnica-Krezel, 2007).

The phenotypes observed in both the somites and endothelium after loss of RA signaling are not the result of a simple deficit of mesoderm. RA signaling loss of function affects somite maturation and morphogenesis, but does not prevent them from forming, and yet the migration of the angioblasts is still disrupted. The direct transcriptional targets of RA signaling that induce somite maturation are unknown, but it is clear that RA signaling controls the timing and progress of somite shape changes as they mature. Cell autonomous effects of RA depletion on the somites, coupled with angioblast migration defects in RA depleted somites, indicate RA directly controls somite shape changes that facilitate migration. The fact that somites can adopt mature shapes in explants, even in absence of adjacent somites, indicates this control is local to individual somites. It is also noteworthy that loss of retinoic acid signaling activity in a relatively few number of adaxially localized cells in our chimeric somites were sufficient to cause somite shape and angioblast migration defects. This implies that the somite cells most medial to the midline could be the population required for the ventral expansion of the somite and NES. It is not clear if NES is required for the initiation of angioblast migration, as some midline-directed cellular movement occurs in DEAB treated embryos before NES is visibly seen. It is possible that the somite itself could facilitate the angioblasts movement to the midline, or simply provide an opening through which angioblasts can reach the midline. The fact that angioblasts reside adjacent to the notochord in tbx16 mutants, but do not migrate beneath them, implies angioblasts are not sufficient to drive NES or their coalescence at the midline. It is likely that ventral expansions of the somite drives the notochord away from the endoderm. Together, this indicates that somitic maturation requires tight spatial and temporal control to facilitate the concomitant development of the endothelium.

## Materials and Methods

### Generation of the tg(hsp70l:lifeact-mscarlet) transgenic line

This transgenic line was generated using the plasmid pNJP002 (hsp70l:lifeact-mScarlet) and tol2 transgenesis (Kawakami, 2004; Kikuta and Kawakami, 2009). pNJP002 was generated using PCR amplification paired with Gibson assembly cloning. A restriction digest was performed on a tol2-hsp70 plasmid (Row et al., 2016) using the restriction enzymes BamHI-HF and ClaI (NEB bio labs). PCR amplification of plasmid hCCR4 (lifeact-mScarlet) was performed using forward primer CAAGCTACTTGTTCTTTTTGCAGGATCCATGGGCGTGGCCGACTTG and reverse primer TTCGTGGCTCCAGAGAATCGATTCACTTGTACAGCTCGTCCATGC. Gibson Assembly (NEBbuilder HiFi DNA Assembly, NEB bio labs) was performed on restriction digested hCCR2 and PCR amplified lifeact-mscarlet to generate plasmid pNJP002. The tg(hsp70l:lifeact-mscarlet) line was generated by injecting 25 picograms of pNJP002 with 25 picograms of tol2 mRNA using methods described previously (Row et al., 2016).

### In-situ hybridization and immunohistochemistry

Whole-mount in situ hybridization was performed as previously described (Griffin et al., 1995). An antisense RNA probe was synthesized as previously described for etv2 (Sumanas et al., 2005). The cldn5b, cxcr4a, apela, aplnra, aplnrb, cxcl12a and dab2 DIG probes were generated by taq polymerase-generated PCR fragments integrated into a pCRII vector using the Topo-TA Cloning kit (Thermo Fisher). Immunohistochemistry was performed by fixing embryos overnight in 4% Paraformaldehyde (Sigma) dissolved in phosphate buffered saline containing 0.1% Tween-20 (PBST) at 4°C. Permeabilization was performed by the placing the embryos in PBS containing 2% Triton X-100 (Sigma) for 1 hour with agitation. Embryos were washed 3x with PBST and blocked in a blocking solution (PBST, 10% goat serum, 1% BSA) for 2 hours at room temperature. Primary antibodies, Etv2 Rabbit Polyclonal and GFP mouse monoclonal, were diluted to 1:500 in blocking solution and incubated overnight at 4°C. Embryos were washed 5x with PBST for 20 mins each wash. Embryos were then incubated in blocking solution for 2 hours; and then incubated in the secondary antibodies Alexa Fluor 488 and Alexa Fluor 568 diluted 1:1000 in blocking solution at 4°C overnight. DAPI stains were done in a 10 µg/mL dilution in PBST for at least 1 hour with agitation.

### Histological analysis

Sections were fixed overnight in 4% Paraformaldehyde (Sigma) dissolved in phosphate buffered saline containing (PBS) at 4°C. Nuclei labeling was done for whole embryos in a 10 µg/mL dilution of DAPI in PBST for at least 1 hour with agitation. Sections, after fixation and DAPI staining, were done in PBST with a 0.15mm microknife (Fine Science Tools), at the indicated somite location for embryos up to 24 hpf. At 24 hpf or older, embryos were sectioned at approximately the 7th somite. Sections were mounted in 1% Agarose in PBS in a in a 35mm glass bottom dish with uncoated #1.5 coverslip and 20mm glass diameter (MatTek).

### Microinjections

A mix of two tbx16 morpholinos (MO1: AGCCTGCATTATTTAGCCTTCTCTA (1.5 ng) and MO2: GATGTCCTCTAAAAGAAAATGTCAG (0.75 ng)) were prepared as previously reported and injected into tg(kdrl:GFP) embryos (Row et al., 2016). 2.5 ng of control morpholino and noto morpholino were injected into tg(kdrl:GFP) embryos (Ouyang et al., 2009). 200 ng of mcherry-caax mRNA was injected into tg(kdrl:GFP) embryos.

### Microscopy and Imaging

DIC and fluorescent time-lapse images of wild-type, noto morphants, aldh1a2 mutants, tbx16 mutants, and drug treated embryos were performed using a Leica DMI6000B inverted microscope. Wild-type and control embryos were siblings of the morphants and mutants. Live embryos were mounted in 1% low-melt agarose in embryo media containing 1x tricaine (25x stock 0.4g/l;Pentair,TRS1) in a 35mm glass bottom dish with uncoated #1.5 coverslip and 20mm glass diameter (MatTek). In-situ hybridization experiments were imaged using a M165FC microscope (Leica) equipped with an Infinity 3 camera (Lumenera). Embryos were mounted in either horizontal or flat mounted configuration in 70% glycerol on glass slides.

Fluorescent images of control, tbx16, and noto morphant (Figure 3) cross sections were imaged on a custom assembled spinning disk confocal microscope consisting of an automated Zeiss frame, a Yokogawa CSU-10 spinning disc, a Ludl stage controlled by a Ludl MAC6000 and an ASI filter turret mated to a Photometrics Prime 95B camera. All other cross sections, including explants, were performed on a custom-assembled spinning disk confocal microscope consisting of a Zeiss Imager A.2 frame, a Borealis modified Yokogawa CSU-10 spinning disc, ASI 150uM piezo stage controlled by an MS2000, and ASI filter wheel, and a Hamamatsu ImageEM x2 EMCCD camera (Hamamatsu C9100-23B). These microscopes were controlled with Metamorph microscope control software (V7.10.2.240 Molecular Devices), and laser power levels were set in Vortran’s Stradus VersaLase 8 software. Images were processed in Fiji.

### Drug treatments

RA was depleted in embryos using N,N-diethylaminobenzaldehyde (DEAB) and BMS453. Stock concentrations of 20 mM DEAB in DMSO were diluted to working concentrations of 20 µM in embryo media. Stock concentrations of 2 mM BMS453 in DMSO were diluted to working concentrations of 2 µM in embryo media. Treatments were done on shield stage embryos unless otherwise noted. All-trans RA was used to activate the RA pathway. Stock concentrations of All-trans RA were diluted from 1 mM stock to 0.1 µM working concentrations in embryo media. All treatments for all-trans RA were performed in tailbud stage.

### Generation of explants for imaging

Transgenic or mutant embryos were grown to the 10 somite stage and then transferred to Modified Barth’s Saline (MBS) (Sigma). In this medium, embryos were anesthetized with 1 x Tricaine (25x stock 0.4g/l; Pentair, TRS1) and had their chorions manually removed by forceps. The embryos were then sectioned with a 0.15mm microknife (Fine Science Tools). Excess yolk was removed with the microknife, leaving some yolk to prevent injury to the endoderm, and the embryo was then transferred to 35 mm glass-bottom dish (Matek) with a fire polished pipette. 1.3 % low gelling temperature agarose in MBS was heated to 40°C and placed over the embryo in the proper orientation. The agarose was allowed to solidify and the explant was time-lapsed using fluorescent and DIC microscopy.

### Quantification and Statistical Analysis

When determining the fluorescence of angioblasts at the midline of the embryo, we utilized the integrated density feature of Fiji software for image analysis. We calculated the corrected total fluorescence of the area at the midline and of the whole embryo, using the notochord as a reference for the midline. Corrected total fluorescence for the midline was calculated as: (mean gray value of midline x area of midline) - (mean gray value of background x area of midline). Corrected total fluorescence for the embryo was calculated: (mean gray value of embryo x area of embryo) - (mean gray value of background x area of embryo). Percentages for fluorescent intensity were calculated as (corrected total fluorescence at the midline) / (corrected total fluorescence of the whole embryo). Tracks for cells were made using the Trackmate plug-in for Fiji (Tinevez et al., 2017). We utilized a Laplacian of Gaussian detector and a Simple LAP Tracker to generate the tracks and analysis. P-values for indicated figures were generated using two-tailed unpaired student t-tests. Graphs were generated using Graphpad Prism 8.4.2 for dot plots, and Excel for line graphs. The black lines indicate the median and interquartile ranges, and were generated using the Prism software. Simple linear regression was also determined using Graphpad software.

## Acknowledgments

We thank David Matus for critical review of the manuscript, as well as members of the Martin and Matus labs for helpful comments. We thank Jesus Torres Vazquez for sending the kdrl transgenic reporter lines. We thank Neal Bhattacharji and Stephanie Flanagan for excellent zebrafish care. We thank Tianying Chen for assistance in data acquisition. We thank Rebecca Adikes, Thom Geer and Nobska Imaging for microscopy support. The illustration of the 15-somite stage zebrafish embryo was created by Biorender.com. This work was supported by the NSF (IOS 1452928) and NIH NIGMS (R01GM124282).

## Supplemental Figures

**Figure S1.**
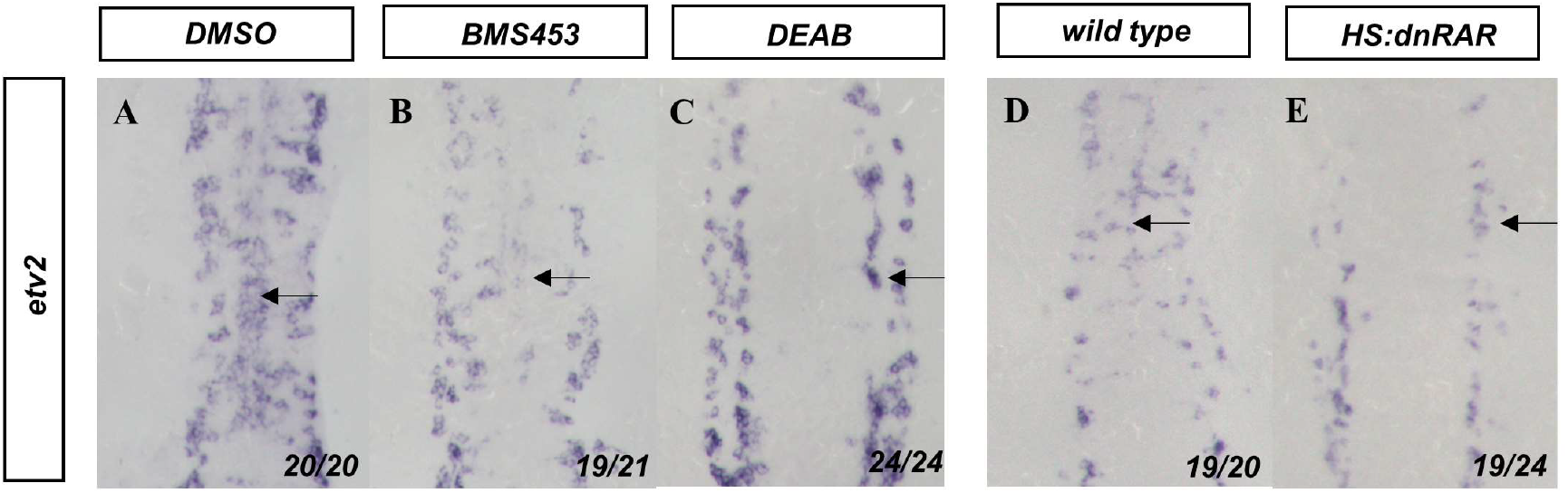
Three alternative methods of RA signaling disruption inhibit midline angioblast migration. Angioblasts are labelled by etv2 in-situ hybridization in 13-somite stage embryos (black arrows) after (A) DMSO, (B) BMS453, or (C) DEAB treatment, or after bud stage heat-shock in (D) wild-type and (E) HS:dnrar embryos. Embryos are shown from a dorsal view with anterior to the top.

**Figure S2.**
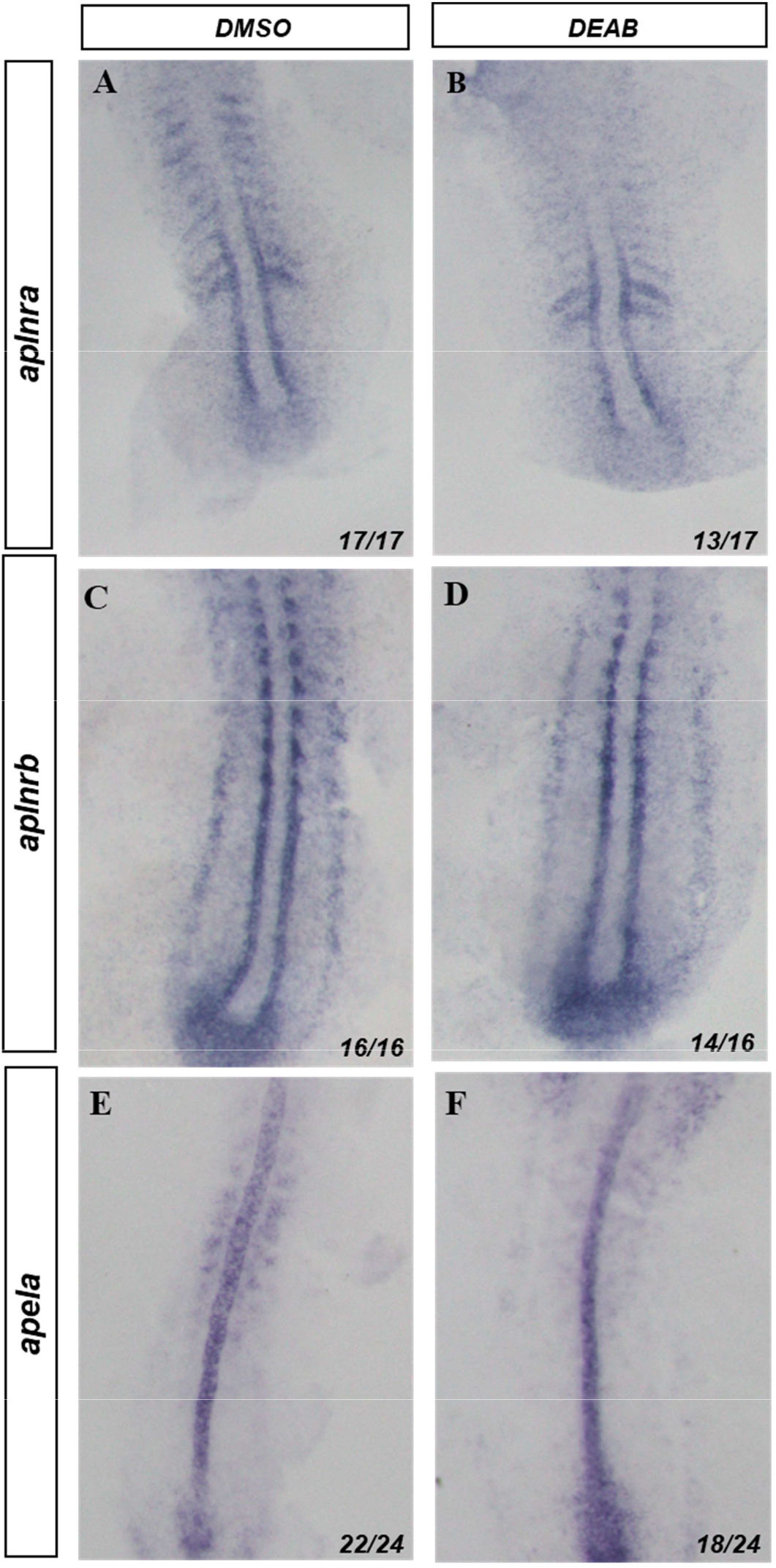
Apela/Aplnr axis is not altered by changes in retinoic acid signaling In situ hybridization of 10-somite stage embryos. (A-B) In-situ hybridization against aplnra. (A) DMSO treated embryos. (B) DEAB treated embryos. (C-D) Insitu hybridization against aplnrb. (C) DMSO treated embryos (D) DEAB treated embryos. (E-F) In-situ hybridization against apela. (E) DMSO treated embryos (F) DEAB treated embryos

**Figure S3.**
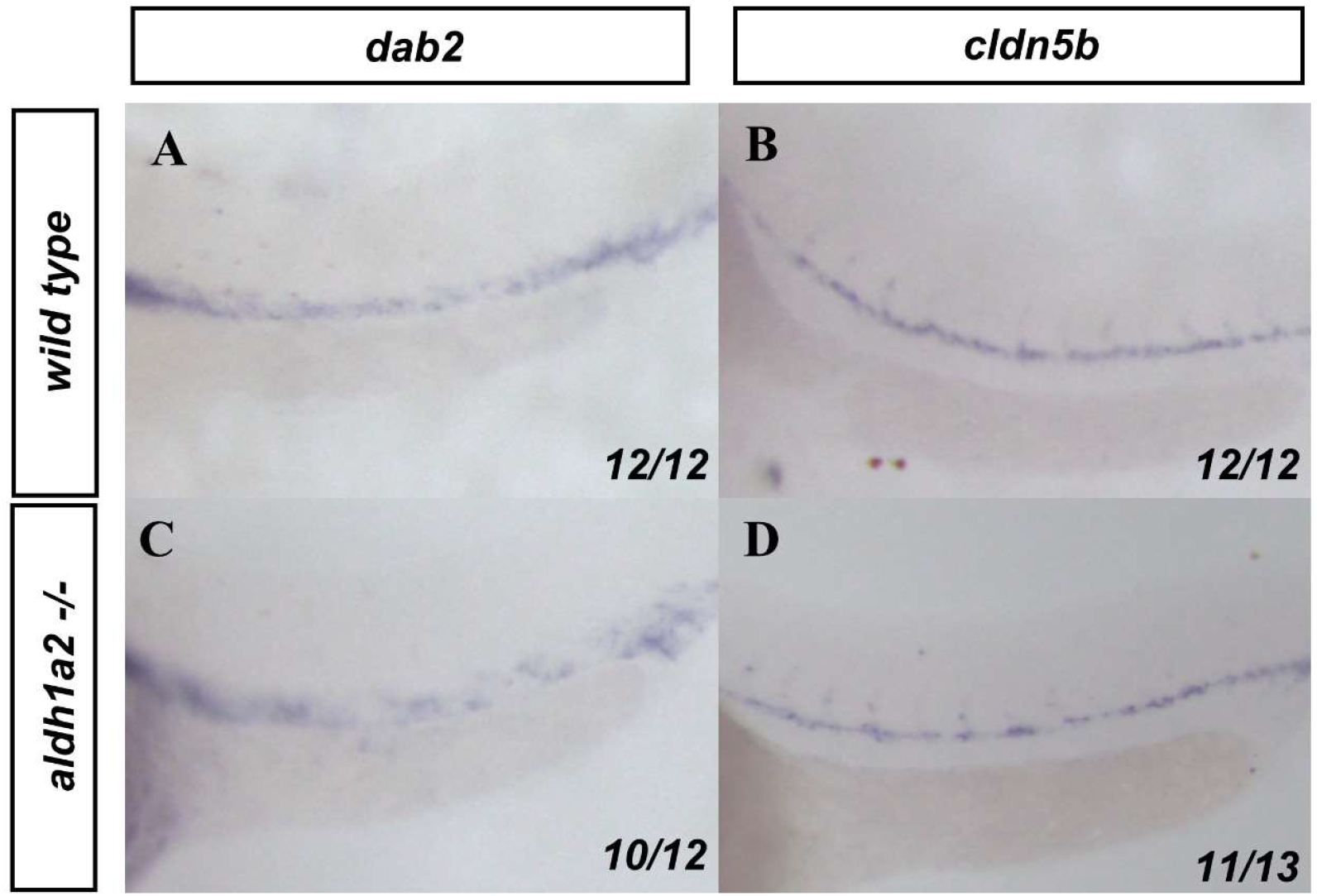
Effects of RA depletion on arterial and venous markers. (A, C) dab2 and (B, D) cldn5b in-situ hybridization labels the veins and arteries, respectively in (A-B) wild-type or (C-D) aldh1a2 -/- embryos. Embryos are shown from a lateral view with anterior to the left.

**Figure S4.**
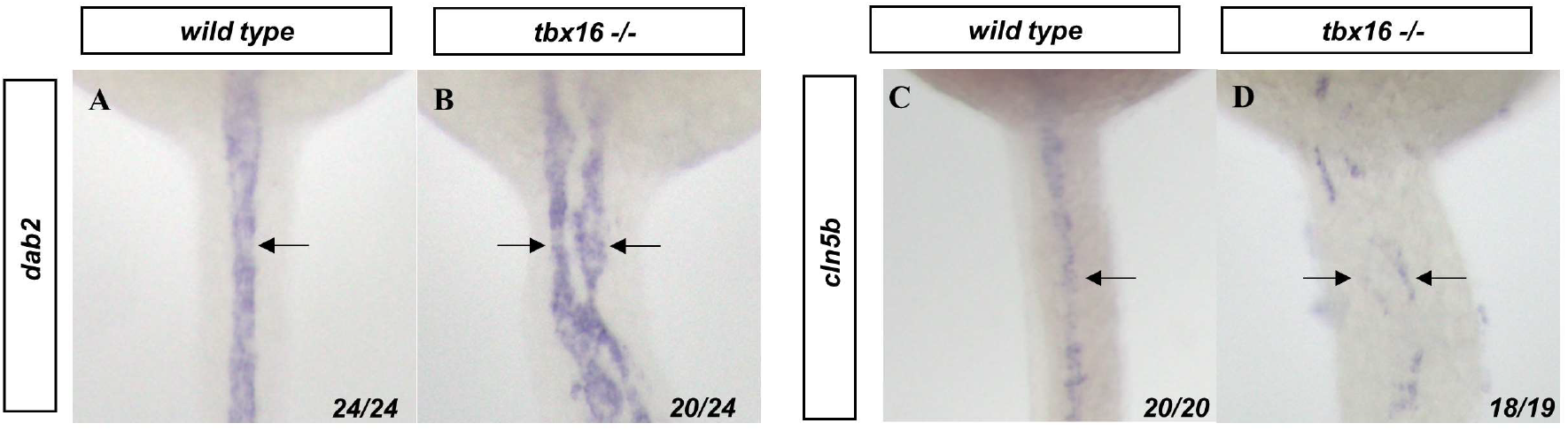
Bifurcated angioblasts in tbx16 mutants preferentially join the venous population. (A, B) dab2 and (C, D) cldn5b in-situ hybridization labels the veins and arteries, respectively (black arrows) in (A,C) wild-type and (B,D) tbx16 -/- embryos. Embryos are shown from a dorsal view with anterior to the top.

**Figure S5.**
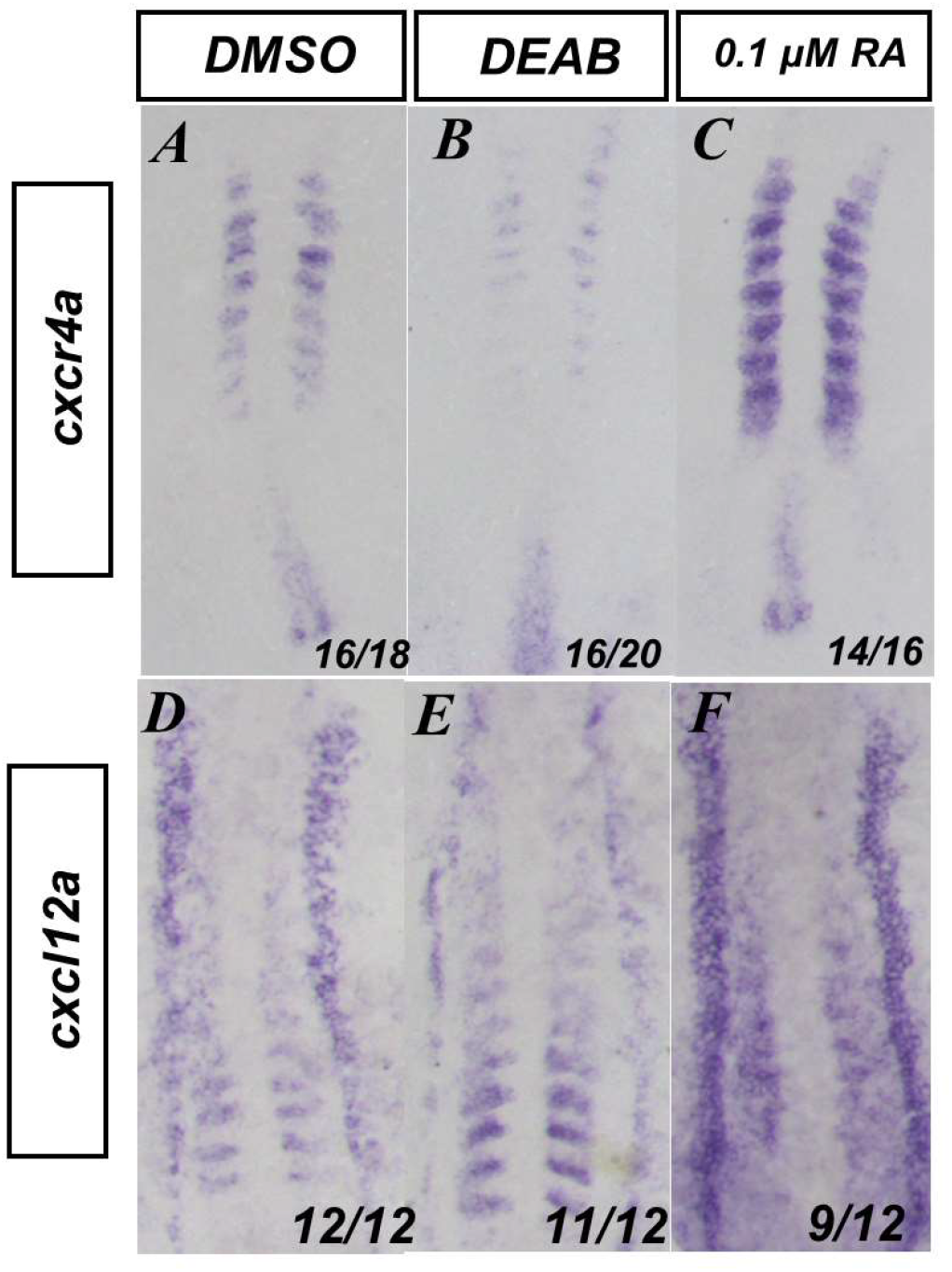
Retinoic acid signaling affects the expression of genes associated with somitic movements. (A-C) In-situ hybridization for cxcr4a and (D-F) cxcl12a in (A, D) DMSO, (B, E) 20 μM DEAB, or (C, F) 0.1 μM RA treated embryos. Embryos are shown from a dorsal view with anterior to the top.

Movie S1. Time-lapse fluorescent imaging of tg(kdrl:eGFP) embryo at the 10 somite-stage. tg(kdrl:eGFP) marks angioblasts as they migrate to the midline. Angioblasts display the anterior posterior processivity while coalescing at the midline. Frame rate= one image/5 min. Run time= 235 min.

Movie S2. Time-lapse fluorescent imaging of tg(kdrl:eGFP), aldh1a2 -/- embryo at the 10 somite-stage. tg(kdrl:eGFP) marks angioblasts as they migrate to the midline. Angioblasts lose anterior to posterior processivity and show disrupted migration. Frame rate= one image/5 min. Run time= 235 min.

Movie S3. Time-lapse fluorescent imaging of a 10 somite-stage tg(kdrl:eGFP) embryo with addition of 0.1 μM RA at the tailbud stage. tg(kdrl:eGFP) marks angioblasts as they migrate to the midline. Angioblasts accelerate their processivity towards the midline. Frame rate= one image/5 min. Run time= 235 min.

Movie S4. Time-lapse fluorescent imaging of HS:id3, HS;dnRAR cells in a tg(kdrl:eGFP) host. Cells expressing HS:id3, HS;dnRAR migrate to the midline along with angioblasts in a tg(kdrl:eGFP) host. Cells with dnRAR migrate faithfully with angioblasts. Frame rate= one image/5 min. Run time= 180 min.

Movie S5. Time-lapse fluorescent imaging of tg(tbxta:kaedae), tg(actc1b:gfp) trunk explant. Trunk explant shows notochord displacement away from ventral-most portion of somites. Frame rate= one image/15 min. Run time= 285 min.

Movie S6. Time-lapse confocal fluorescent imaging of a HS:mcherry-caax-p2a- NLS-kikume embryo in DMSO treatment. HS:mcherry-caax-p2a-NLS-kikume marks nuclei within the somite. Sample tracks ranging from red (most displacement) to blue (least displacement) show broad movement within the somite. Frame rate= one image/5 min. Run time= 240 min.

Movie S7. Time-lapse confocal fluorescent imaging of a HS:mcherry-caax-p2a- NLS-kikume embryo in DEAB treatment. HS:mcherry-caax-p2a-NLS-kikume marks nuclei within the somite. Sample tracks ranging from red (most displacement) to blue (least displacement) show little directional movement in 20 μM DEAB treatment conditions. Frame rate= one image/5 min. Run time= 240 min.

Movie S8. Time-lapse fluorescent movie of mcherry-caax mRNA injected tg(kdrl:eGFP) explant at the 10 somite-stage. tg(kdrl:eGFP) marks angioblasts as they migrate to the midline and mcherry-caax marks the cell surfaces. A trunk explant, imaged around the 5th somite, shows normal migration of the angioblasts to the midline. Frame rate= one image/5 min. Run time= 280 min.

Movie S9. Time-lapse fluorescent movie of mcherry-caax mRNA injected tg(kdrl:eGFP) explant at the 10 somite-stage, treated with 20 μM DEAB. tg(kdrl:eGFP) marks angioblasts as they migrate to the midline. The trunk explant shows angioblast migration defects seen in fixed DEAB sections. Frame rate= one image/5 min. Run time= 175 min.

## Notes

### Competing Interest Statement

The authors have declared no competing interest.

